# Transinfection of *Wolbachia w*AlbB into *Culex quinquefasciatus* mosquitoes does not alter vector competence for Hawaiian avian malaria (*Plasmodium relictum* GRW4)

**DOI:** 10.1101/2024.02.16.580617

**Authors:** A. Marm Kilpatrick, Christa M. Seidl, Isaiah J. Ipsaro, Chris E. Garrison, Giulia Fabbri, Paul I. Howell, Austin G. McGowan, Bradley J. White, Sara N. Mitchell

## Abstract

Avian malaria is expanding upslope with warmer temperatures and driving multiple species of Hawaiian birds towards extinction. Methods to reduce malaria transmission are urgently needed to prevent further declines. Releasing *Wolbachia*-infected incompatible male mosquitoes suppress mosquito populations and releasing *Wolbachia*-infected female mosquitoes could reduce pathogen transmission if the *Wolbachia* strain reduced vector competence. We cleared *Culex quinquefasciatus* of their natural *Wolbachia pipientis w*Pip infection and transinfected them with *Wolbachia w*AlbB isolated from *Aedes albopictus*. We show that *w*AlbB infection was transmitted transovarially, and demonstrate cytoplasmic incompatibility with wild-type mosquitoes infected with *w*Pip from Oahu and Maui, Hawaii. We measured vector competence for avian malaria, *Plasmodium relictum*, lineage GRW4, of seven mosquito lines (two with *w*AlbB; three with natural *w*Pip infection, and two cleared of *Wolbachia* infection) by allowing them to feed on canaries infected with recently collected field isolates of Hawaiian *P. relictum*. We tested 73 groups (N_total_ = 1176) of mosquitoes for *P. relictum* infection in abdomens and disseminated (thorax) infections 6-14 days after feeding across a range of parasitemias from 0.028% to 2.49%, and a smaller subset of salivary glands. We found no measurable effect of *Wolbachia* on any endpoint, but strong effects of parasitemia, days post feeding, and mosquito strain on both abdomen infection prevalence and disseminated infection prevalence. These results suggest that releasing male *w*AlbB-infected *C. quinquefasciatus* mosquitoes could suppress *w*Pip-infected mosquito populations, but would have little positive or negative impact on mosquito vector competence for *P. relictum* if *w*AlbB became established in local mosquito populations. More broadly, the lack of *Wolbachia* effects on vector competence we observed highlights the variable impacts of both native and transfected *Wolbachia* infections in mosquitoes.

## Introduction

Hawaiian birds are experiencing an extinction crisis. Of the 47 species of Hawaiian honeycreepers known to science, forty-one are extinct or federally endangered (Paxton *et al*. 2022). One of the key threats to the remaining species is avian malaria, *Plasmodium relictum* (lineage GRW4), transmitted by *Culex quinquefasciatus* mosquitoes (Beadell *et al*. 2006; Paxton *et al*. 2022). Rising global temperatures due to climate change have increased the distribution of mosquito populations and the transmission of malaria in higher elevations on multiple islands, as predicted two decades ago (Benning *et al*. 2002). This has led to further population declines in many species, with two additional species now nearly extinct in the wild (Atkinson *et al*. 2014; Paxton *et al*. 2022). New tools are urgently needed to reduce transmission of avian malaria.

One recent breakthrough for reducing transmission of mosquito-borne diseases is the use of *Wolbachia* bacteria either to suppress mosquito populations or to reduce mosquito vector competence (Flores & O’Neill 2018; Ross *et al*. 2019). *Wolbachia* bacteria are intracellular parasites that naturally infect many mosquitoes and insect species (Hilgenboecker *et al*. 2008), and they can be transinfected into new species or populations (Hughes & Rasgon 2014). Different strains of *Wolbachia* can have different effects on mosquitoes, including reducing adult lifespan, reduced vector competence for some pathogens, and a reduction in mosquito populations through cytoplasmic incompatibility (CI) (Iturbe□Ormaetxe *et al*. 2011). CI caused by *Wolbachia* results in inviable embryos unless both male and female mosquitoes have compatible *Wolbachia* strains. Control strategies involving the release of *Wolbachia*-infected incompatible males could be further improved if the *Wolbachia* strain also reduces vector competence in accidentally released female mosquitoes. Similarly, if a *Wolbachia* strain reduces vector competence, then population replacement via large scale releases of female mosquitoes infected with this *Wolbachia* strain could reduce vector competence and transmission, as has been recently demonstrated for dengue virus and *Aedes aegypti* mosquitoes (Utarini *et al*. 2021; Velez *et al*. 2023). The EPA recently approved an emergency exemption to release large numbers of *w*AlbB transinfected male mosquitoes to reduce populations of *C. quinquefasciatus* in Hawaii (US EPA, 2023), making this a viable strategy to reduce transmission of avian malaria in Hawaii.

Our goals were threefold: first, to transinfect *Wolbachia w*AlbB into a *C. quinquefasciatus* line to use for incompatible male releases for population suppression in Hawaii; second, to confirm complete maternal transmission of *Wolbachia* infection and CI of transinfected males with wild-type females from Hawaii infected with *Wolbachia w*Pip; and third, to determine if the transfinfected *Wolbachia w*AlbB strain alters the vector competence of transinfected female *C. quinquefasciatus* mosquitoes for the avian malaria lineage found in Hawaii, *P. relictum* GRW4. We did not have an *a priori* hypothesis about whether the *w*AlbB strain of *Wolbachia* would increase or reduce vector competence of *C. quinquefasciatus* for *P. relictum* because effects of transinfected *Wolbachia* in other mosquito species have been highly variable (Hughes *et al*. 2014; Ross *et al*. 2019) and no studies of the effects of transinfected *Wolbachia* on malaria competence have been done in *C. quinquefasciatus*. Previous studies of *C. quinquefasciatus* naturally infected with *Wolbachia w*Pip found higher susceptibility for *P. relictum* lineage SGS1 than mosquitoes cleared of *w*Pip with antibiotics (Zélé *et al*. 2014a). However, there was no effect of *Wolbachia w*Pip on infection prevalence in the field (Zélé *et al*. 2014b). Finally, a previous study that created a *w*AlbB transinfected *C. quinquefasciatus* line didn’t quantify the effects on vector competence for any pathogen (Ant *et al*. 2020), and this line has been lost in an insectary malfunction (Steven Sinkins, personal communication).

## Methods

### *Culex quinquefasciatus* Mosquito Strains

We studied eight *C. quinquefasciatus* mosquito lines (Table S1) which were combinations of four mosquito strains (Palmyra Atoll (abbreviated Palmyra or Palm in figures), Oahu, Maui, Field (Captain Cook, Hawaii)) with one of two *Wolbachia* strains (native *w*Pip or transinfected *w*AlbB) or cleared of *Wolbachia* infection via antibiotic treatment (“None”). We refer to lines using the strain and *Wolbachia* type (e.g. Palmyra-*w*Pip, or Oahu-None). We used subsets of these eight lines for three types of experiments: maternal inheritance of transinfected *w*AlbB *Wolbachia*, cytoplasmic incompatibility, and vector competence for avian malaria, *P. relictum* GRW4 (Table S1; Supplemental Material, Methods: Mosquito strains).

### Clearing *Wolbachia w*Pip and transinfecting *Wolbachia w*Alb

We created two mosquito lines without *Wolbachia*, Palmyra-None and Oahu-None, by clearing native *Wolbachia* wPip with antibiotics and then used these to create two additional mosquito lines transinfected with *Wolbachia* wAlbB (Supplementary material: Methods: Transinfection of *Culex quinquefasciatus* with *Wolbachia w*AlbB). We examined maternal transmission of *w*AlbB in Palmyra-*w*AlbB (“DQB”, *D*ebug *q*uinquefasciatus wAlb*B*.”), and bi-directional cytoplasmic incompatibility between Palmyra-*w*AlbB and both Oahu-*w*Pip and Maui-*w*Pip (Supplementary material, Methods: Maternal transmission determination and Cytoplasmic incompatibility (CI) testing).

### *Plasmodium relictum* isolates

We collected two isolates of *Plasmodium relictum* (lineage GRW4) from wild birds at two sites on Hawai’i Island, one from an ‘Apapane (*Himatione sanguinea*) from Pu’u Wa’awa’a Forest Reserve (19.738154°N, 155.875234°W, 1,230 m above sea level) in February 2020 and another from a Warbling White-eye (*Zosterops japonicus*) from the same Captain Cook, HI site where Field mosquitoes were collected (Supplemental Material: Methods: *Plasmodium relictum* isolates).

### Experimental *Plasmodium relictum* infections and mosquito feeding

We inoculated canaries intramuscularly with 50-200 µL of whole blood (parasitemia: 0.1– 2.85%) containing an avian malaria isolate that had been passaged 1–7 times and was either a thawed deglycerolized sample, or was fresh blood from another infected canary. Starting on day 5 post-infection afterward we took 5-10 µL of blood by brachial venipuncture and screened thin blood smears by microscopy (Supplemental Material: Methods: *Plasmodium relictum* isolates) and by qPCR (Neddermeyer *et al*. 2023; Paxton *et al*. 2023) to detect infection and estimate parasitemia. Once infection was detected, we allowed approximately 50 mosquitoes from each of three mosquito lines to feed on each infected bird simultaneously. We differentiated the three mosquito lines during feedings by spraying each line with a green or red fluorescent marker or leaving it unmarked (Figure S1; Supplementary Material, Methods: Mosquito marking); the line receiving each spray color (or none) was randomly selected. We fed most mosquitoes by restraining an infected canary on top of a container containing the mosquitoes that had holes in the lid that allowed the bird’s legs and feet to be inside the container (Figure S2a). Field mosquitoes were more hesitant to feed and were placed in a Bugdorm with an unrestrained bird in a PVC cylinder with a perch and allowed to feed for ∼ 8 hrs overnight (Figure S2b). We collected engorged mosquitoes with an aspirator and transferred them into a cage in an incubator set to 26 °C. We provided mosquitoes with cottons soaked in a 10% sucrose solution and held them until dissection. We collected unfed females, knocked them down on ice, and counted the number of each spray color under a UV light to quantify feeding success for each group.

We killed engorged mosquitoes 6-14 days after feeding and dissected them by cleanly separating the thorax from the abdomen at the scutellum with sterile dissection needles. We placed abdomens in 96-well DNA extraction plates containing Chemagic DNA lysis buffer (PerkinElmer, Waltham, MA, USA) and placed the head, thorax, and legs of each individual in a second plate to test for a disseminated infection. When DNA extraction plates were not available, abdomens and thoraxes (including the head and legs) were placed in separate 1 mL vials containing 0.5 mL of 70% ethanol. All samples were stored at -20 °C until DNA extraction and processing by qPCR (Supplemental Information, Methods: *Plasmodium relictum* qPCR and ddPCR). We also examined *P. relictum* infection in the salivary glands for a subset of 106 Oahu mosquitoes of all three Wolbachia types (*w*AlbB, *w*Pip, and None) (Supplemental Information, Methods: Salivary gland infection).

### Statistical analyses

We analyzed the fraction of mosquitoes testing positive for *P. relictum* by qPCR with generalized linear models with a binomial distribution and a log link. We analyzed abdomen and thorax infections separately. We included log(Parasitemia), days post feeding, mosquito strain (Oahu, Palmyra, or Field), and *Wolbachia* strain (*w*AlbB, *w*Pip, or None) as predictors. We excluded small batches of mosquitoes with N<4 (a batch is a unique mosquito strain+*Wolbachia* strain+day post feeding+parasitemia), but results were qualitatively identical if we included all mosquitoes. We examined two and three-way interactions among the predictors log_10_(Parasitemia), days post feeding, and mosquito strain and used AIC to determine which interactions were best supported by the data. We did not include spray color or *P. relictum* isolate in the final model because neither had support when added to the best fitting model (see Table S4; Spray color: □^2^ = 1.47, df = 2, P = 0.48; PUWA Isolate coef.: -0.225, SE = 0.18, Z = - 1.23, P = 0.22).

To examine salivary gland infection, we log transformed the ratio of *P. relictum* DNA to mosquito DNA in the salivary glands and analyzed it with a linear model with *Wolbachia* type as a predictor. There was heteroscedasticy in the residuals so we performed a robust comparison using the *coeftest* function in the *lmtest* package with a variance-covariance matrix estimated using the *sandwich* package. All analyses were performed in R, v.4.3.1.

## Results

### Transinfected line maternal transmission and Cytoplasmic Incompatibility (CI)

We cleared *w*Pip *Wolbachia* infection from the Palmyra-*w*Pip strain of *C. quinquefasciatus* with antibiotics and successfully transinfected a single female with the *w*AlbB strain of *Wolbachia*. We then examined maternal transmission in 991 adults from generations 7 thru 13 of the mosquito line from this female (N_ave_ = 188.8 (range 187-192)/generation, excluding generation 9 which had 32 mosquitoes). Every sample from every generation tested positive for *w*AlbB by ddPCR, indicating complete maternal transmission of *Wolbachia* across generations (mean ratio of wAlbB gene copies to *C. quinquefasciatus* gene copies: 2.95; SE = 0.077).

Bidirectional cytoplasmic incompatibility between mosquitoes infected with *w*AlbB and *w*Pip was nearly complete in both directions. In crosses between Palmyra-*w*AlbB males (“DQB”) with Oahu-*w*Pip females, only 0.93% of 18,457 eggs hatched (99.07% were inviable), and with Maui-*w*Pip females, only 1.54% of 13,898 eggs hatched (98.46% were inviable) (Figure 1; Table S2). Similarly, when Palmyra-*w*AlbB females were mated with Oahu-*w*Pip males only 0.03% of 10,496 eggs hatched (99.97% inviable), and when crossed with Maui-wPip males only 0.05% of 14,706 eggs hatched (99.95% inviable) (Figure 1; Table S3).

**Figure 1.**
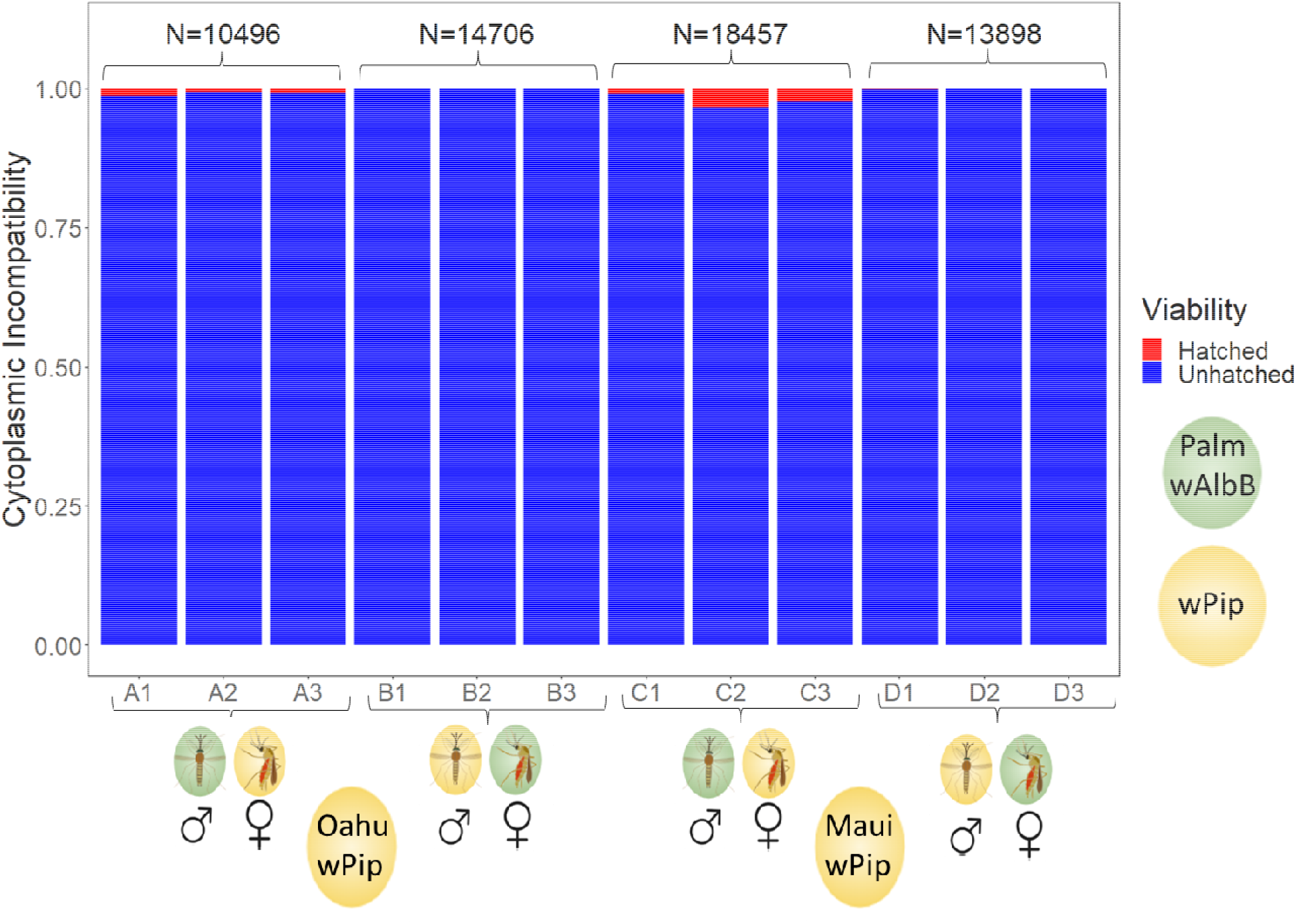
Bi-directional cytoplasmic incompatibility results between Palmyra mosquitoes infected with *Wolbachia wAlbB* (green ovals) and two mosquito strains (Oahu and Maui) infected with native *Wolbachia wPip* (yellow ovals). Three replicates are shown for each of four crosses that include both males and females of each mosquito strain: A1-A3: Palmyra-*w*AlbB males X Oahu-*w*Pip females; B1-B3: Oahu-*w*Pip males X Palmyra-*w*AlbB females; C1-C3: Palmyra-*w*AlbB males X Maui-*w*Pip females; D1-D3: Maui-*w*Pip males X Palmyra-*w*AlbB females. Each of the 12 replicates had N_mean_ = 4796 (range 786-7131); the total number of eggs for each cross is shown above the bars.

### Vector competence measurements

We infected fourteen canaries with *P. relictum* GRW4, which had peak parasitemias averaging 1.22% (range 0.04% - 4.7%; note that we didn’t sample birds every day and could have missed individual peak parasitemias). Peak parasitemia increased with passage number (Log(peak parasitemia) = -6.81+ 0.465 (SE 0.19) ^*^ Passage number; P = 0.032; N= 13; R^2^ = 35.3%), but was unrelated to the dose injected (# of parasites), or isolate (dose coeff.: -0.39, SE 4.02, N = 13, P = 0.925; isolate coef., PUWA vs CACO: -0.042, SE 0.68, N = 13, P = 0.952, in separate models with passage number). Over the course of infection, birds had daily parasitemias of 0.016% to 4.7% and we used parasitemias between 0.028% and 2.49% to measure vector competence of *C. quinquefasciatus* mosquitoes (Figure S3).

We fed 68 batches (mean N = 57; SD = 17.9; range 20-130) of *C. quinquefasciatus* mosquitoes on these infected canaries (Figure S4; Table S4; see Supplementary Material: Feeding success). We tested the 1176 fed mosquitoes from these 68 batches in 73 groups for abdomen and thorax infection, by qPCR (mean N = 16/group; SD = 7.5; range 4-46). Despite this enormous number of groups, there was no evidence for differences in disseminated (thorax) infection prevalence among *Wolbachia* strains (*w*AlbB, native *w*Pip, or None) (compare the height of points with the same color in each panel of Figure 2 and height of different colored lines in Figure 3; Table S5).

**Figure 2.**
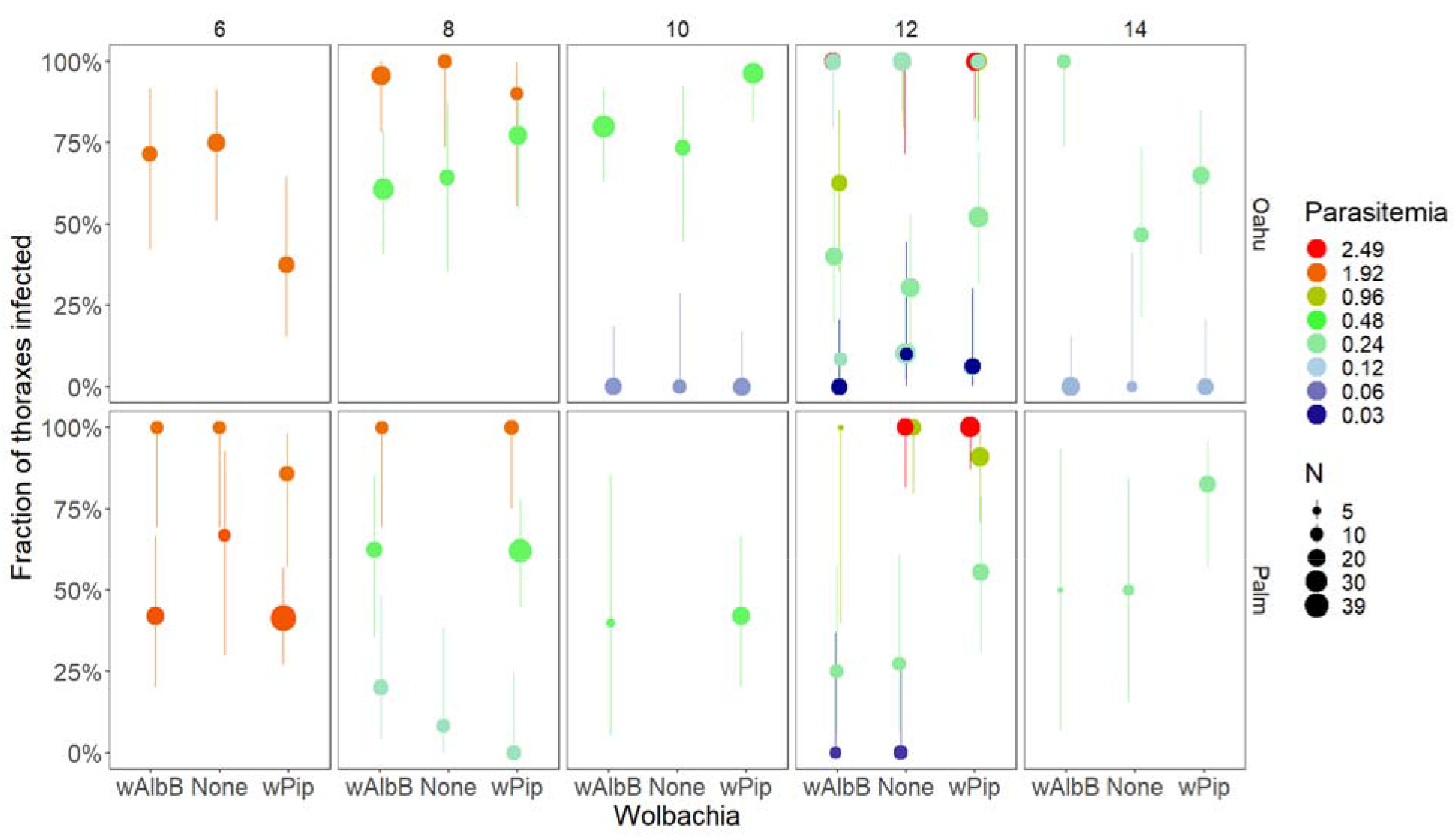
Fraction of thoraxes infected (and binomial 95% CI) plotted against *Wolbachia* type in the mosquitoes (*w*AlbB, *w*Pip, or None). The color shows the parasitemia (percent of red blood cells infected) of the bird the mosquitoes fed upon (on a log_2_ scale), the panel rows show the mosquito strain (Oahu or Palmyra), the panel columns show the days post-feeding when the mosquitoes were dissected (6-14 days), and the size of the points shows the sample size for each point (range 4-39). The fitted model (Table S5) indicates there are no consistent differences among *Wolbachia* types (points of the same color in each panel are, on average, at the same height). Points have been slightly jittered along the x-axis to aid in visualization. Note that in a small number of experiments one of the three *Wolbachia* groups had insufficient mosquitoes that successfully fed and is not shown (e.g. bottom middle panel, Palm-None on day 10).

**Figure 3.**
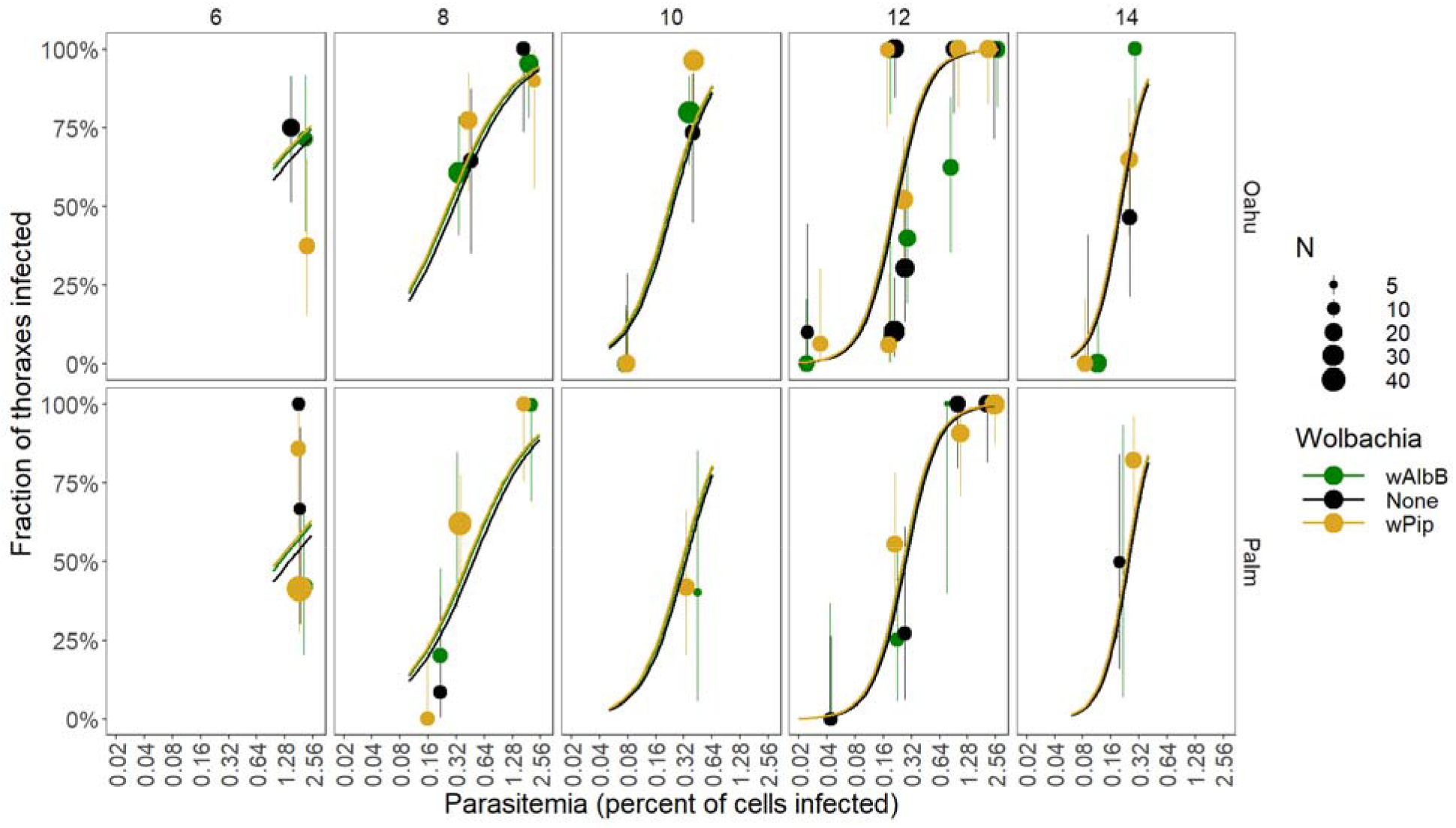
Fraction of thoraxes infected (and binomial 95% CI) plotted against the parasitemia (percent of red blood cells infected) of the bird the mosquitoes fed upon (on a log_2_ scale). This is the same data as in Figure 2, plotted with a different x-axis. Color shows the *Wolbachia* type in the mosquitoes (*w*AlbB, *w*Pip, or None), the panel rows show the mosquito strain (Oahu or Palmyra), the panel columns show the days post-feeding when the mosquitoes were dissected (6-14 days), and the size of the points shows the sample size (range 4-39). The lines show the fitted model for each of the three *Wolbachia* types, which are on top of each other and difficult to distinguish because there was no measurable differences among *Wolbachia* types (Table S5). Points have been slightly jittered along the x-axis to aid in visualization.

However, the fraction of mosquitoes with disseminated infections increased sharply with parasitemia and days since feeding, and the slope of parasitemia increased over time (Figures 2,3, Table S5). Disseminated infection prevalence was slightly higher in the Oahu strain than in the Palmyra strain (Figures S5; Table S5), and highest in the Field strain (Figure S5; Table S5). The Results were qualitatively identical when we analyzed each *Wolbachia*-mosquito strain pair separately (Figure S6; Table S6).

Abdomen infection prevalence was higher than disseminated infection prevalence, but patterns were similar, with there being no detectable effect of *Wolbachia* type, and infection prevalence increased with parasitemia, and differed between the mosquito strains (Figures 4, S7; Table S7). Abdomen infection prevalence increased faster with days since infection for the Palmyra and Field strains than for the Oahu strain, which resulted in differences between the mosquito strains changing with days since feeding (prevalence was higher in Oahu mosquitoes 6-10 days after infection but was higher in the Palmyra and Field strains 12-14 days after infection; Figures S7; Table S7).

**Figure 4.**
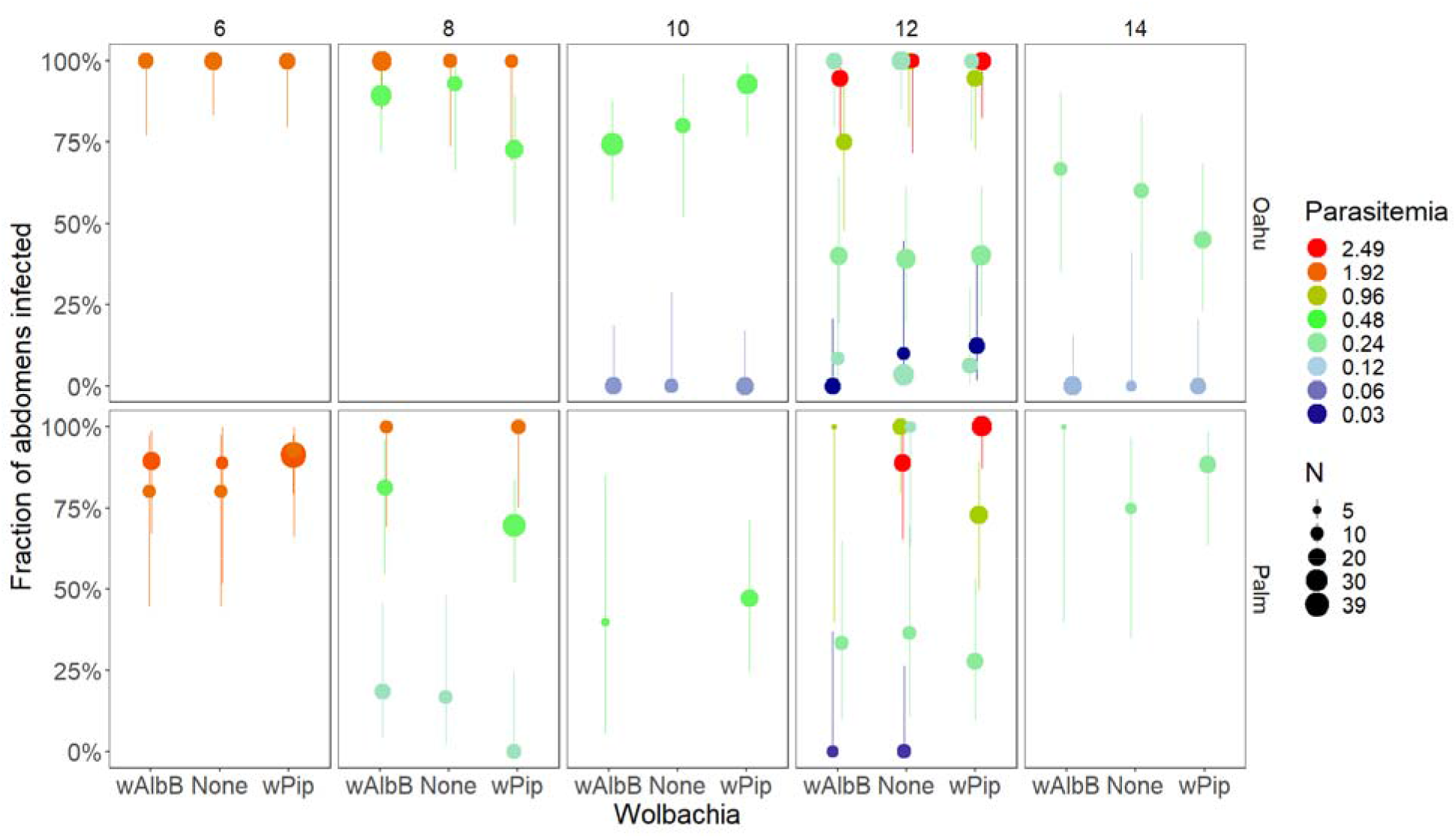
Fraction of abdomens infected (and 95% CI) plotted against *Wolbachia* type in the mosquitoes (*w*AlbB, *w*Pip, or None). The color shows the parasitemia (percent of red blood cells infected) of the bird the mosquitoes fed upon (on a log_2_ scale), the panel rows show the mosquito strain (Oahu or Palmyra), the panel columns show the days post-feeding when the mosquitoes were dissected (6-14 days), and the size of the points shows the sample size (range 4-39). Points have been slightly jittered along the x-axis to aid in visualization.

We tested salivary glands from 106 Oahu mosquitoes for *P. relictum* DNA by ddPCR that had fed on a canary with a parasitemia of 0.12% 14 days earlier. Salivary glands from all 106 mosquitoes tested positive for *P. relictum* DNA, and there were no differences in the amount of *P. relictum* DNA among the three *Wolbachia* types (Figure 5).

**Figure 5.**
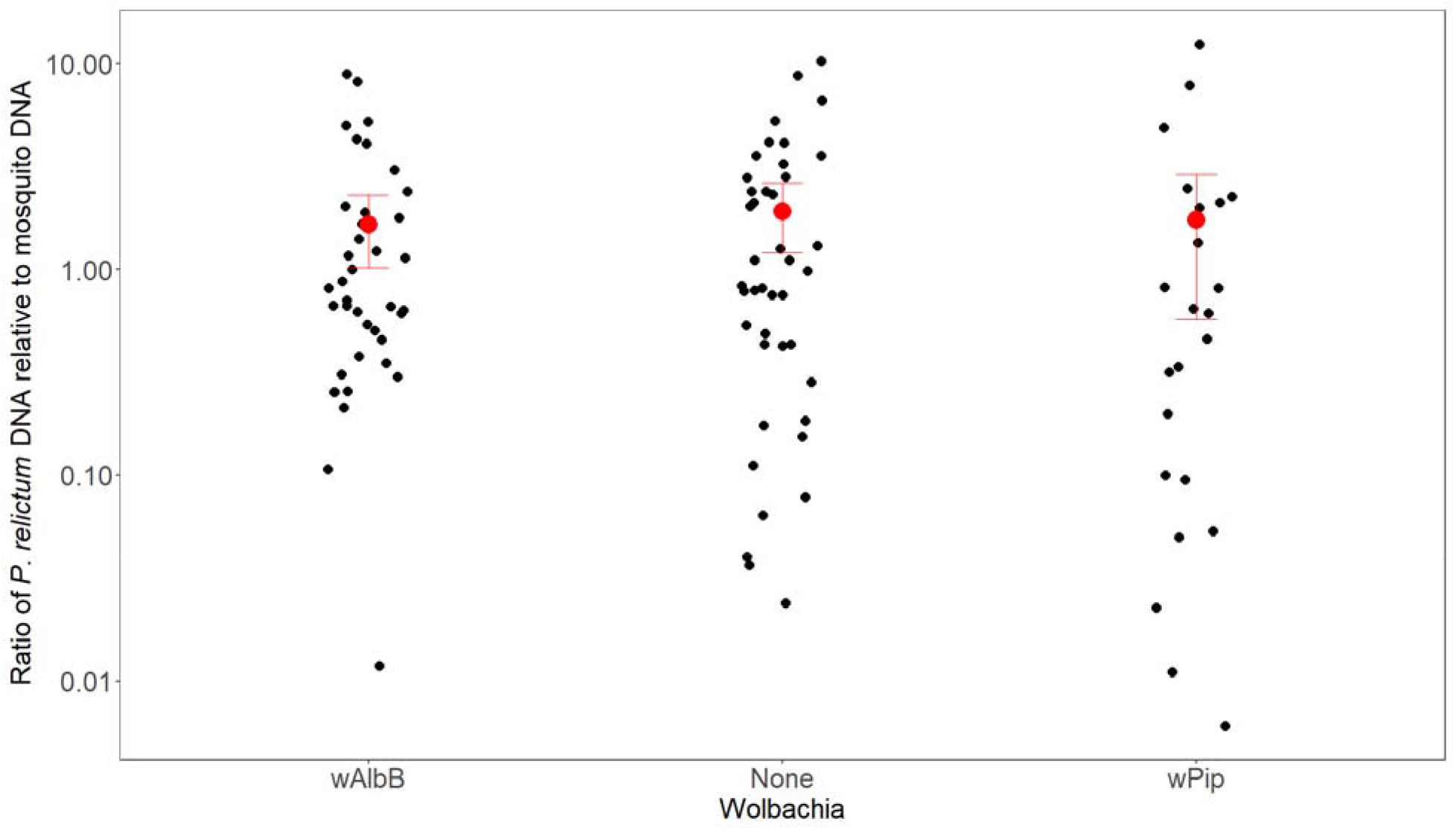
*Wolbachia* strain and salivary gland infection. *P. relictum* DNA measured by ddPCR, on a log scale, in salivary glands from 106 Oahu mosquitoes infected *w*AlbB, *w*Pip, or no *Wolbachia* infection. Red points show mean and 95% CI. There was no difference among *Wolbachia* types (robust F = 1.16, df = 2, P = 0.32).

## Discussion

We successfully transinfected the *w*AlbB *Wolbachia* strain into *C. quinquefasciatus* mosquitoes originating from Palmyra Atoll after clearing them of their natural *w*Pip *Wolbachia* infection. The resultant Palmyra-*w*AlbB line of mosquitoes exhibited complete maternal transmission of *w*AlbB across generations, and almost 100% bidirectional CI with mosquitoes from Hawaii (Maui and Oahu) infected with *w*Pip. We then created another *w*AlbB-infected line by backcrossing the Palmyra-*w*AlbB into an Oahu strain of *C. quinquefasciatus* that we cleared of natural *w*Pip infection. The creation of these two lines of *w*AlbB-infected *C. quinquefasciatus* mosquitoes makes population suppression of *C. quinquefasciatus* via widespread release of *w*AlbB-infected males possible both in Hawaii and elsewhere. The bidirectional CI we observed between *w*AlbB and *w*Pip mosquitoes would result in any accidentally released *w*AlbB-infected females having inviable offspring if they mate with native *w*Pip males.

We then examined the effect of *w*AlbB, *w*Pip or no *Wolbachia* on vector competence in two mosquito strains, Palmyra and Oahu, for *P. relictum* GRW4, the only lineage of avian malaria in Hawaii (Beadell *et al*. 2006). We found infection with *Wolbachia* strains *w*AlbB and *w*Pip had no detectable effect on vector competence of *C. quinquefasciatus* mosquitoes for avian malaria *P. relictum* GRW4. There were no detectable differences among *Wolbachia* groups in thorax infection or abdomen infection in either the Oahu or Palmyra mosquito strains. We also found no difference among *Wolbachia* groups in the amount of *P. relictum* DNA in salivary glands in a subset of mosquitoes from the Oahu strain. This is the first study, to our knowledge, to assess the impact of both natural and stable transinfections of *Wolbachia* on malaria vector competence in a natural mosquito-pathogen system. Taken together these results indicate that the purposeful or accidental introduction of the *w*AlbB strain of *Wolbachia* to mosquito populations in Hawaii would neither help nor hurt conservation efforts to reduce transmission of avian malaria (Wild 2023).

Our results differ from two previous studies that found that *Wolbachia* altered *Plasmodium* prevalence in malaria vectors. In *C. quinquefasciatus*, natural *w*Pip infections increased vector competence for *P. relictum* lineage SGS1 compared to those cleared of *Wolbachia* infection (Zélé *et al*. 2014a). In contrast, in *Anopheles stephensi, w*AlbB reduced midgut infection prevalence of *Plasmodium falciparum* (Bian *et al*. 2013). The conflicting results between these two studies suggests that the impacts of *Wolbachia* on *Plasmodium* infection are not uniform, which is consistent with the highly variable effects of transinfected *Wolbachia* on other pathogens in other vectors (Ross *et al*. 2019). We had much larger sample sizes than all past studies combined, and examined seven lines of mosquitoes, including two inbred mosquito strains with two strains of *Wolbachia* or no *Wolbachia* and one Field mosquito strain, suggesting that a lack of an effect of *Wolbachia* on *Plasmodium* vector competence was not due to a lack of power.

In contrast to the lack of differences among *Wolbachia* lines, we found large differences among mosquito strains in vector competence, with Field mosquitoes having the highest vector competence, followed by the inbred Oahu strain and then the inbred Palmyra strain. The differences in vector competence among mosquito strains indicates that we had ample power to detect differences in vector competence. They also underline the extensive variability in vector competence among populations of the same mosquito species, which is common for many vector-pathogen pairs (Kilpatrick *et al*. 2010; Reisen *et al*. 2008).

We found a relatively steep relationship between parasitemia and both thorax and abdomen infection, and the relationship became steeper with time since feeding. Twelve days after feeding the relationship was very steep, with parasitemias up to 0.1% leading to relatively few disseminated thorax infections (<15.0%), whereas parasitemias only three-fold higher (0.3%) led to most (73.0%) mosquitoes having disseminated infections. In contrast, eight days after feeding, a 10-fold range of parasitemias (0.06% to 0.6%) produced a similar range in thorax infection. The steep relationship between host parasitemia and disseminated infection prevalence made it more challenging to determine the appropriate day to feed mosquitoes on a canary and which day to dissect and test mosquitoes to obtain an intermediate level of infection. In many of our feedings most or almost none of the mosquitoes in all groups had disseminated infections (Figures 2, 3, S5). This steep relationship, if it is also present in wild mosquitoes, would lead to highly heterogeneous infectiousness among birds (and bird species), with birds infecting nearly all or almost none of the mosquitoes that fed on them depending on whether their parasitemia was above or below a relatively narrow threshold parasitemia (e.g. ∼0.1%-0.3% on day 12 post feeding). Our data from the Field mosquito strain was limited, with only one day post feeding (day 12) where we had fed mosquitoes on canaries with a range of parasitemias (four parasitemias, ranging from 0.028% to 1.225%; Figure S7). Analysis of just this (small) dataset of wild G0 mosquitoes produced a much less steep relationship (Figure S7), suggesting the steep relationship we observed for the dataset composed of mosquitoes from two inbred lines (Oahu and Palmyra), may be a result of limited genetic variation.

In summary, the rapid decline of many species of Hawaiian birds over the last decade, due to a climate-change driven increase in malaria transmission at higher elevations, requires urgent action to prevent extinction. We created two lines of *w*AlbB-infected *C. quinquefasciatus* mosquitoes with high CI with Hawaiian *C. quinquefasciatus* infected with wPip, which could be used to suppress mosquito populations via widespread release of *w*AlbB-infected males. We examined whether the transinfected *Wolbachia* strain *w*AlbB increased or decreased vector competence of these two lines for *P. relictum* GRW4. We found no effect of *Wolbachia* infection on thorax, abdomen, or salivary gland infection, suggesting that replacement of the current *Wolbachia* in Hawaiian mosquitoes (*w*Pip) with the *w*AlbB strain would have little impact on their susceptibility to infection and ability to transmit this parasite. Furthermore, vector competence in one population of mosquitoes collected directly from the field had much higher infection rates than the two transinfected lines. This suggests that accidental releases of small numbers of female mosquitoes with *Wolbachia* strain *w*AlbB with the large numbers of male mosquitoes with *Wolbachia* strain *w*AlbB that is planned for 2024 (Wild 2023) is unlikely to alter the transmission dynamics of malaria beyond the effect of the males greatly reducing mosquito populations.

## Acknowledgements

We thank Nancy McInerney, Nichelle VanTassel, and Rob Fleischer for genetic analysis of our thawed *P. relictum* isolate to confirm it was lineage GRW4. Funding was provided by NSF grants DEB-1717498 and DEB-1911853.

## Notes

### Competing Interest Statement

GF, PIH, AGM, BJW, and SNM report employment and equity ownership at Verily Life Sciences, a for-profit company developing new technologies for mosquito control.

### Summary of Updates

I have added the supplemental material file which was not included in the automatic biorxiv posting by the journal

