## Supplemental materials for "Transinfection of *Wolbachia w*AlbB into *Culex quinquefasciatus* mosquitoes does not alter vector competence for Hawaiian avian malaria (*Plasmodium relictum* GRW4)"

**Methods**

*Culex quinquefasciatus mosquito strains*

We worked with eight mosquito lines which were combinations of mosquito strains and *Wolbachia* strains (Table S1). The Palmyra-*w*Pip strain was started from egg rafts collected from Palmyra Atoll (5.889481°N, 162.078570°W, 5 m above sea level) in 2018 and reared for 40 generations in the laboratory. We created a second line (Palmyra-None) by clearing the Palmyra-*w*Pip line of *Wolbachia* for 5 generations using antibiotics (see below). We created a third line by transinfecting the Palmyra-None line with *wAlbB* (Palmyra-*w*AlbB) (see below).

The Oahu-*w*Pip strain was started from eggs collected from Kawainui Marsh (21.393988°N,  157.752154°W, 1 m above sea level) in 2020 and reared for 10 generations. We created the Oahu-None line by clearing the Oahu-wPip line for 5 generations. We created the Oahu-*w*AlbB line by backcrossing the Oahu-None line with the Palmyra-wAlbB line for seven generations.

The Maui-*w*Pip strain was started using egg rafts collected from Makawao, HI (20.851838°, -156.313887°, 499 m above sea level) in 2020. Finally, the Field-*w*Pip line was started using egg rafts collected from Captain Cook, Hawaii (19.461187°N, 155.896432°W, 204m above sea level) in March 2023; for this line, larvae were reared and G0 individuals were used for measuring vector competence (Table S1).

All 7 colonized lines (all but Field-*w*Pip) were reared using standard protocols (Crawford *et al.* 2020), with 8-10 egg rafts seeded into each 1 Liter rearing pan; adults were fed on avian blood (Lampire); and females were allowed to lay eggs directly on RO water. All mosquito stages were maintained at 26-28 °C in 70-80% humidity. We reared Field-wPip larvae in plastic pans (44 cm × 25 cm × 10 cm) with 200–350 larvae/pan, filled with 1 Liter of deionized water and fed daily 0.2–0.4 g of ground fish food (Koi's Choice® Premium Fish Food). We transferred pupae to 30 $cm^3$ mosquito cages (BugDorm). Emerging adults were fed *ad libitum* on 10%

sucrose solution-soaked cottons. We used 7 to 14-day-old Field-*w*Pip adults and deprived them of sucrose cottons for 24-48 hrs prior to blood feeding in vector competence experiments.

Transinfection of *Culex quinquefasciatus* with *Wolbachia*

We first cleared native *Wolbachia w*Pip infection from the Palmyra-*w*Pip line by exposing adult mosquitoes to a solution of 25mg/100ml tetracycline (Sigma T7660), pH buffered with TrisBase to 7.4, and provided in 10% sugar feeders for five generations (Dobson & Rattanadechakul 2001). We then isolated *Wolbachia w*AlbB from *Aedes albopictus* originating from Kuala Lumpur, Malaysia (KLP strain). We dissected 30-60 ovaries of female KLP mosquitoes and harvested *w*AlbB *Wolbachia* into phosphate buffered saline (PBS) for each transfection experiment. Ovary tissue was thoroughly homogenized using a Tapered Tissue Grinder (DWK Life Sciences Wheaton™) before centrifugation to remove cellular debris. We filtered and centrifuged the supernatant (Pall Acrodisc®) to select for *Wolbachia* cells. We resuspended the pellet in PBS to a concentration of 1-6 ovaries/µL for microinjection into *Culex* embryos.

We then transinfected mosquitoes from the Palmyra-*None* line with wAlbB. We allowed Palmyra-None females to lay eggs on reverse osmosis (RO) water and aligned eggs individually against nitrocellulose membrane (MF-Milipore #HAWP04700) underneath moistened filter paper on a glass slide. We then injected *w*AlbB *Wolbachia* extract into each egg at the posterior end using a beveled quartz glass pipette (Sutter Instruments #QF100-70-10) and the XenoWorks® Digital Microinjector and Micromanipulator system (Sutter Instruments).

Post injection, we stored *w*AlbB eggs in a plastic container with a constant RO water supply and incubated overnight at 28°C before hatching the next day. We hatched injected eggs in a petri dish with RO water, yeast slurry and Koi fish pellets. We maintained flooded eggs under insectary conditions (28°C, 80% humidity, 12:12 hr light cycle) for ~3 days before larvae were counted. We reared larvae to adults on a diet of cricket powder and Koi fish pellets.

We harvested adult males from G0 (transfected eggs) and screened them for the presence of *Wolbachia* using ddPCR (Crawford *et al.* 2020) to detect transfection. We mated females from injection sets with positive G0 males with Palmyra-None males and provided them with a warmed avian blood meal (Lampire) in plastic 100 x 15 mm petri-dishes (VWR, #25384-324) with Parafilm M membrane (Bevis, #HS234526A). Isofemale lines were then established from single blood fed females placed into iso-tubes (Drosophila vials [Genesee Scientific 32-116]) with RO water (~10 mL). Once eggs were laid, females were harvested and individually tested for the presence of *Wolbachia* via ddPCR. If an egg-laying female was positive for *Wolbachia*, its G1 larvae were reared and the process of male and female screening repeated until a stable line with 100% maternal transmission was achieved.

Maternal transmission determination

The Palmyra-*w*AlbB line was allowed to stabilize through sib-mating and standard larva-to-adult rearing for six generations before maternal transmission was assessed. We then quantified maternal transmission of *w*AlbB *Wolbachia* in at least 95 adult males and 95 females in each of five generations (G7, 8, 10, 11, 12; G9 had insufficient samples). We extracted mosquito DNA and quantified *w*AlbB presence using ddPCR on the *WSP* gene (Crawford *et al.* 2020) with the following modifications: we used the following for the ddPCR primer probe set for the *C. quinquefasciatus* housekeeping gene *RPL5* (ensembl ID LOC6043019):  F 5' GGCAACACTGACATCTCTGT 3', R 5' AAAATCGAAGATGGCGTTGC 3', probe 5' HEX CTCGCTCGCTCTCGGCTCTCCT ZEN/3' IBFQ. A few samples (15/991) were omitted from analysis because the mosquito reference gene *RPL5* was less than 100 droplets, which indicated poor DNA quality/yield.

Cytoplasmic incompatibility (CI) testing

We assessed CI in the Palmyra-*w*AlbB transinfected line by mating Palmyra-*w*AlbB males to two lines with native *w*Pip infections, Oahu-*w*Pip and Maui-*w*Pip (Table S1). We also measured bidirectional CI by mating Oahu-*w*Pip and Maui-*w*Pip males with Palmyra-*w*AlbB females. For each mating assay, pupae were microscopically assessed for sex before transfer to eclosion cages (BugDorm-4S3030, W30 x D30 x H30cm) with provision of 10% sucrose *ad libitum.* When sexually mature (day four post eclosion) 150 males from one of two *Wolbachia* strains (*w*AlbB or *w*Pip) were mated with 100 females of the other *Wolbachia* strain (*w*Pip or *w*AlbB, respectively) in medium cages (BugDorm-4S3030, W30 x D30 x H30cm) for three nights. After mating, females were provided an avian blood meal (Lampire) in a plastic 100 x 15mm petri-dish (VWR, #25384-324) with Parafilm M membrane (Bevis, #HS234526A). Following blood feeding, females were allowed to oviposit individually in iso-tubes (Genesee Scientific 32-116) with RO water (~10 mL) or *en masse* in oviposition bowls (Eco-Products, EP-BSC8-GS) with ~ 100 mL of RO water. Rafts were collected and eggs counted under the microscope before hatching, then returned to RO water to hatch for seven days before any larvae were counted. For each of the four mating crosses (Palmyra-*w*AlbB males with female Oahu-*w*Pip and Maui-*w*Pip; and Palmyra-*w*AlbB females with male Oahu-*w*Pip and Maui-*w*Pip) three biological replicates were performed.

<u>*Plasmodium relictum* isolates</u>

We collected two isolates of *Plasmodium relictum* (lineage GRW4) from wild birds at two sites on Hawai'i Island, one from an ‘Apapane (*Himatione sanguinea*) from Pu’u Wa’awa’a Forest Reserve (19.738154°N, 155.875234°W, 1,230 m above sea level) in February 2020 and another from a Warbling White-eye (*Zosterops japonicus*) from the same Captain Cook, HI site where mosquitoes were collected. We took 50-100 µl of blood from the brachial vein of birds and placed it into a 1 mL syringe with an appropriate volume of citrate–phosphate–dextrose solution with adenine (CDPA; Sigma-Aldrich, St. Louis, MO, USA) to create a 1:9 CDPA to blood ratio (Carlson *et al.* 2016) and stored it up to 48 hrs at 4 °C. We used 1–2 µl of blood from the same bird to create a thin blood smear that was air-dried for 30 minutes, fixed in 100% methanol, and later stained for one hour with Giemsa (Valkiunas 2004). We screened each stained smear by examining 50 microscope fields at 1000x magnification using oil immersion to determine the presence and parasitemia of a sample. If we observed one or more cells infected with malaria, we inoculated the blood-CDPA mixture intramuscularly into the pectoral muscle of a domestic canary (*Serinus canaria*). We passaged isolates one to seven times in canaries via intramuscular inoculations of 50-100 µl of infected blood between birds before exposure to mosquitoes in feeding trials, or before cryopreservation in glycerolyte for future use (Moll *et al.* 2013). Both isolates used in mosquito feedings had at least 1 passage between the wild source bird and the bird used for mosquito feedings. All work with wild and laboratory birds was performed under animal care and use protocol Kilpm2003 approved by the Institutional Animal Care and Use Committee at the University of California in Santa Cruz, USA.

<u>Mosquito Marking</u>

In order to differentiate between mosquito lines (*Wolbachia* strains) during the feeding assay, mosquito lines were randomly chosen to either be unmarked or sprayed with a green or red fluorescent marker (SmartWater CSI LLC, GBR) (Figure S1). The two markers used were Cartax-DP Fluorescent Marker (green) and Invisible Red S marker (red), which were combined with a polymer to ensure adherence to the mosquitoes (Faiman *et al.* 2021). For the green marker, we mixed Cartax-DP Fluorescent Marker with Mowilith LDM 7709 Polymer *in house* using methods modified from (Faiman *et al.* 2021). Briefly, 60 µL of Cartax-DP Fluorescent Marker (0.5% sodium lauryl sulfate [SLS] as dispersal agent) was combined with 12 µL of Mowilith polymer in 3,928 µL of water to produce a 1.5% dye solution with a dye:polymer ratio of 5:1. To aid visualization of the red marker, we used a premixed dye-polymer also available from SmartWater (SmartTrace, SmartWater Forensic Marker Solution), which is a 0.5% Invisible Red

S dye solution containing a Invisible Red SMarker:Mowilith LDM 7709 Polymer ratio of 1:1 (0.5% SLS). The fluorescent mixtures (*in house* or premixed) were then vortexed for 15 to 30 seconds and added to a nebulizer (ASOMI Mesh Nebulizer) reservoir. The nebulizer mouthpiece was connected to a plastic container containing 150 female mosquitoes and turned to the highest setting for thirty seconds. During spraying the container was tapped lightly to ensure mosquito movement for maximum coverage. The spray was allowed to dry for two hours and a small subset of mosquitoes were checked for fluorescent marking under a UV light to confirm proper application. We randomly rotated the marking of mosquito strains (red, green or no spray) for each feeding experiment.

The accuracy of this marking method was tested by molecularly identifying 95 mosquitoes that we had identified using the color marking (20 Palmyra-None marked red; 16 Palmyra-*w*AlbB marked green; 20 Oahu-None marked red; 32 Oahu-*w*Pip (unmarked); and 7 Oahu-*w*AlbB marked green). We used ddPCR to examine the presence/absence of *w*Pip or wAlbB *Wolbachia* (see Crawford *et al.* 2020) with wPip *WSP* (ID WP0937) (F 5' GCTGGTGCTCGTTATTTTGG 3', R 5' ACAGCGCTGTAAAGGACATT 3', probe 5' FAM AAGAAGCAGTATCAGCTACTAAAGAGA ZEN/3' IBFQ). We used the RPL5 *C. quinquefasciatus* housekeeping gene as a positive ddPCR control. All but one mosquito (94/95) was correctly identified using the marking colors: one unmarked Oahu-*w*Pip mosquito was mis-identified as a red marked Oahu-None.

<u>*Plasmodium relictum* qPCR and ddPCR</u>

We extracted DNA from all mosquito samples for qPCR and ddPCR analysis using the Chemagic 360 DNA (PerkinElmer) extraction system (see Crawford *et al.* 2020 for methods), and analyzed for the GRW4 lineage of *P. relictum*. For real-time qPCR, the following oligos were used: Forward primer - 5' ATTAGCAGAACAAAGAAACTTAACA 3'; Reverse primer - 5' CATAGAATGAACATATAAACCAG 3' (Zehtindjiev *et al.* 2008); and Probe - 5'

GCTTTTGGTGCAAGAGAGTATTCAGT 3’ (Whitaker, Kelsey 2022). The qPCR reaction mixture and set up of thermocycler profile follow the methods developed in (Videvall *et al.* 2021). Samples were considered positive for *P. relictum* GRW4 if they crossed the threshold at or before 40 cycles.

We diluted a subset of extracted DNA from salivary glands 1:20 with water and then analyzed it using a QX200 AutoDG Droplet Digital PCR (ddPCR) System (Bio-Rad, Laboratories). The ddPCR is a multiplexed reaction targeting both the *P. relictum GRW4* gene (Forward: 5’ AAATGAGTTTCTGGGGTGCT 3’, Reverse: 5’ TGGGTCACTTACAAGATATCCAC 3’, Probe: 5’ FAM CACATATCCATGAAACTAGTCCAGG3’) and the *Culex quinquefasciatus* RPL5 housekeeping gene (as detailed previously). The housekeeping gene was used to ensure quality of extracted DNA and normalize titer levels between mosquito samples. Thermal cycling conditions were 95°C for 10 minutes, followed by 40 cycles of 95°C for 30 second denaturing step and an annealing step at 55°C for 30 seconds. This was followed by a final extension at 98°C for 10 minutes. Analysis was performed using the QuantaSoft Analysis Pro Software (Bio-Rad).

<u>Salivary Gland Infection</u>

We dissected one or both salivary lobes from each mosquito and washed the lobe(s) three times in phosphate buffer solution to ensure that no stray tissues were still attached to the glands. We placed glands into a deep well plate with 200uL of Lysis buffer and then homogenized them with two 3mm Grinding Balls (OPS Diagnostics) at 900 RPM for two minutes. We placed the plate into a VWR Ultrasonic Cleaner and sonicated it at 35kHz for ten minutes to ensure salivary tissues were completely lysed. We then extracted DNA using the Chemagic 360 DNA (PerkinElmer) extraction system and ran DNA on the QX200 AutoDG Droplet Digital PCR System (Supplemental Information, Methods*: Plasmodium relictum* qPCR and ddPCR).

### Results

<u>Feeding success</u>

Feeding success, or the fraction of mosquitoes that obtained a blood meal, varied both among mosquito strain and *Wolbachia* type (Figure S5; Table S4). Oahu mosquitoes were more successful at feeding than the Palmyra strain and all three Oahu *Wolbachia* groups fed similarly (Figure S5; Table S4). Palmyra mosquitoes with *Wolbachia w*Pip had higher feeding success than Palmyra mosquitoes with the wAlbB *Wolbachia* strain or those cleared of *Wolbachia* (Figure S5; Table S4). In addition, somewhat surprisingly, mosquitoes sprayed red had significantly lower feeding success than those sprayed green or with no spray, which did not differ significantly from each other (Figure S5; Table S4). This difference may be due to a higher polymer  concentration in the red premix dye vs the green markers, or some other factor.

**Supplemental Tables and Figures**

**Table S1. *Culex quinquefasciatus* mosquito lines used for three types of experiments (VC - vector competence for avian malaria (*Plasmodium relictum* GRW4), MI - maternal inheritance of *Wolbachia* strain *w*AlbB, CI - Cytoplasmic incompatibility).** Mosquito lines are referred to using their mosquito strain and *Wolbachia* strain joined with a dash (e.g. Palm-*w*AlbB). wPip clade assignment was performed using molecular markers Ank2 and PK1 (Dumas *et al.* 2013).

| **Line** | **Origin** | **Wolbachia** | **Experiments** | **Notes** |
|---|---|---|---|---|
| Palm-*w*Pip | Palmyra Atoll, USA | *w*Pip (clade 3) | VC | Colony in generation since 2018 |
| Palm-None | Palmyra Atoll, USA | None (*w*Pip cleared) | VC | Created from Palmyra-*w*Pip |
| Palm-*w*AlbB | Palmyra Atoll, USA | *w*AlbB (KLP) transinfected | VC, MI, CI | Original *w*AlbB line; referred to as DQB3 in government documents |
| Oahu-*w*Pip | Oahu, USA | *w*Pip (clade 5) | VC, CI | Colony in generation since 2020 |
| Oahu-None | Oahu, USA | None (*w*Pip cleared) | VC | Created from Oahu-*w*Pip |
| Oahu-*w*AlbB | Oahu, USA | *w*AlbB (KLP) infected via backcross | VC | Generated by crossing males from line Oahu-None with females from line Palm-*w*AlbB for 7 generations |
| Maui-*w*Pip | Maui, USA | *w*Pip | CI | Colony in generation since 2020 |

| | | | | |
|---|---|---|---|---|
| Field | Hawaii, USA | *w*Pip | VC | G0 wild mosquitoes, collected 2023 |

**Table S2. Cytoplasmic Incompatibility (CI) results for the Palmyra-*w*AlbB line for three biological replicates.** Each replicate consisted of 150 males of the Palmyra-*w*AlbB line mated to 100 females infected with *Wolbachia w*Pip from Oahu or Maui. Eggs were assessed for hatching either from individual females (Isofemales) or from eggs laid *en masse* in cages (Mass Cages).

| Replicate | Male | Female | Number of eggs | Number of larvae | Percent CI conferred |
|---|---|---|---|---|---|
| 1 | Palmyra-*w*AlbB | Oahu-wPip | 6264 | 89 | 98.58% |
| 2 | Palmyra-*w*AlbB | Oahu-wPip | 5062 | 31 | 99.39% |
| 3 | Palmyra-*w*AlbB | Oahu-wPip | 7131 | 52 | 99.27% |
| | | **Oahu total** | **18457** | **172** | **99.07%** |
| 1 | Palmyra-*w*AlbB | Maui-wPip | 7914 | 71 | 99.10% |
| 2 | Palmyra-*w*AlbB | Maui-wPip | 786 | 26 | 96.69% |
| 3 | Palmyra-*w*AlbB | Maui-wPip | 5198 | 117 | 97.75% |
| | | **Maui total** | **13898** | **214** | **98.46%** |
| | | **Totals** | **32355** | **386** | **98.81%** |

**Table S3. Cytoplasmic Incompatibility results for female Palmyra-wAlbB line with males of two wPip lines for three biological replicates.** Each replicate consists of 150 males from Oahu or Maui infected with *w*Pip mated to 100 Palmyra-*w*AlbB females. Eggs were assessed for hatching either from individual females (Isofemales) or from eggs laid *en masse* in cages (Mass Cages).

| Replicate | Male | ***Wolbachia*** **female** | **Number of eggs** | **Number of larvae** | **Percent CI conferred** |
|---|---|---|---|---|---|
| 1 | Oahu | Palmyra-*w*AlbB | 2708 | 0 | 100.00% |
| 2 | Oahu | Palmyra-*w*AlbB | 1401 | 0 | 100.00% |
| 3 | Oahu | Palmyra-*w*AlbB | 6387 | 3 | 99.95% |
| | | **Oahu total** | **10496** | **3** | **99.97%** |
| 1 | Maui | Palmyra-*w*AlbB | 7002 | 5 | 99.93% |
| 2 | Maui | Palmyra-*w*AlbB | 4061 | 2 | 99.95% |
| 3 | Maui | Palmyra-*w*AlbB | 3643 | 0 | 100.00% |
| | | **Maui total** | **14706** | **7** | **99.95%** |
| | | **Total** | **25202** | **10** | **99.96%** |

**Table S4. Statistical analysis of feeding success shown in Figure S4.** The generalized linear mixed effects model with a binomial distribution and a logit link had mosquito strain (reference level: Oahu), *Wolbachia* strain (reference level: *w*AlbB), spray color, a two-way interaction between mosquito strain and *Wolbachia* strain, and a random effect for bird ID as predictors. There was significant variation among spray colors ($\chi^2$ = 28.5, df = 2, P = $6.50x10^{-07}$), mosquito strains ($\chi^2$ = 7.9, df = 1, P = 0.0049), *Wolbachia* types ($\chi^2$ = 20.9, df = 2, P = $2.95x10^{-05}$), and the two-way interaction between mosquito strain and *Wolbachia* strain was also significant ($\chi^2$ = 31.9, df = 2, P = $1.19x^{-07}$). The bird ID random effect variance was 0.50.

| **Predictor** | **Estimate** | **SE** | **z value** | **P-value** |
|---|---|---|---|---|
| **(Intercept)** | 0.42 | 0.24 | 1.73 | 0.084 |
| **Spray(None)** | -0.031 | 0.085 | -0.37 | 0.71 |
| **Spray(Red)** | -0.40 | 0.084 | -4.78 | $1.72x10^{-6}$ |
| **strain(Palm)** | -0.66 | 0.12 | -5.59 | $2.26x10^{-8}$ |
| ***Wolbachia*(None)** | -0.065 | 0.11 | -0.58 | 0.56 |
| ***Wolbachia*(*w*Pip)** | -0.036 | 0.11 | -0.32 | 0.75 |
| **strain(Palm)*Wolbachia(None)** | 0.47 | 0.16 | 2.84 | 0.0045 |
| **strain(Palm)*Wolbachia(wPip)** | 0.94 | 0.17 | 5.65 | $1.63x10^{-8}$ |

**Table S5. Statistical analysis of disseminated (thorax) infections shown in Figures 2, 3, and S5.** The generalized linear model with a binomial distribution and a logit link had log(Parasitemia), days post feeding, mosquito strain (reference level: Oahu), *Wolbachia* type (reference level: *w*AlbB), and a two-way interaction between log(Parasitemia) and days post feeding as predictors. There was significant variation among mosquito strains ($\chi^2 = 102.0$, df = 2, $P < 2x10^{-16}$), but not *Wolbachia* strains ($\chi^2 = 1.15$, df = 2, P = 0.56). This model fit better than a model without the two way log(Parasitemia)*days since feeding interaction (ΔAIC = 81.0), and better than models with additional two-way interactions (Parasitemia*strain ΔAIC = 2.99; Parasitemia*strain+days since feeding*strain; ΔAIC = 2.75)

| Predictor | Estimate | SE | z value | P-value |
|---|---|---|---|---|
| **(Intercept)** | -2.91 | 0.46 | -6.38 | $1.72x10^{-10}$ |
| **log(Parasitemia)** | -1.23 | 0.33 | -3.73 | 0.000193 |
| **Days since feeding** | 0.57 | 0.0579 | 9.84 | $< 2x10^{-16}$ |
| **strain(Palm)** | -0.60 | 0.16 | -3.69 | 0.000227 |
| **strain(Field)** | 2.99 | 0.45 | 6.62 | $3.69x10^{-11}$ |
| ***Wolbachia*(None)** | -0.14 | 0.19 | -0.74 | 0.46 |
| ***Wolbachia*(*w*Pip)** | 0.052 | 0.17 | 0.30 | 0.76 |
| **log(Parasitemia)* Days post feeding** | 0.31 | 0.038 | 8.12 | $4.54x10^{-16}$ |

**Table S6. Statistical analysis of thorax infections using individual mosquito-*Wolbachia* strain pairs shown in Figure S6.** The generalized linear model with a binomial distribution and a logit link had log(Parasitemia), days post feeding, mosquito strain - *Wolbachia* strain pair (reference level: Oahu-*w*AlbB), and a two-way interaction between log(Parasitemia) and days post feeding as predictors. There was significant variation among mosquito strain - *Wolbachia* pairs ($\chi^2 = 114.8$, df = 6, $P < 2x10^{-16}$), but not *Wolbachia* strains within mosquito strains (see coefficients and SEs below).

| **Predictor** | **Estimate** | **SE** | **z value** | **P-value** |
|---|---|---|---|---|
| **(Intercept)** | -2.94 | 0.47 | -6.31 | $2.8x10^{-10}$ |
| **log(Parasitemia)** | -1.23 | 0.33 | -3.73 | 0.00020 |
| **Days post feeding** | 0.57 | 0.058 | 9.85 | $< 2x10^{-16}$ |
| **strain(Oahu- none)** | -0.083 | 0.24 | -0.35 | 0.73 |
| **strain(Oahu-*w*Pip)** | 0.072 | 0.23 | 0.31 | 0.76 |
| **strain(Palm-*w*AlbB)** | -0.53 | 0.28 | -1.93 | 0.053 |
| **strain(Palm-none)** | -0.79 | 0.30 | -2.60 | 0.0093 |
| **strain(Palm-*w*Pip)** | -0.52 | 0.23 | -2.25 | 0.025 |
| **strain(Field-wPip)** | 3.07 | 0.46 | 6.63 | $3.45x10^{-11}$ |
| **log(Parasitemia)* Days post feeding** | 0.31 | 0.038 | 8.12 | $4.58x10^{-16}$ |

**Table S7. Statistical analysis of abdomen infections shown in Figure 4.** The generalized linear model with a binomial distribution and a logit link had log(Parasitemia), days post feeding, mosquito strain (reference level: Oahu), *Wolbachia* strain (reference level: *w*AlbB), and a two-way interaction between days post feeding and mosquito strain as predictors. There was significant variation among mosquito strains ($\chi^2 = 95.9$, df = 2, P < $2 \times 10^{-16}$), but not *Wolbachia* strains ($\chi^2 = 1.27$, df = 2, P = 0.53).

| **Predictor** | **Estimate** | **SE** | **z value** | **P-value** |
|---|---|---|---|---|
| **(Intercept)** | 4.53 | 0.67 | 6.74 | $1.63 \times 10^{-11}$ |
| **log(Parasitemia)** | 1.84 | 0.12 | 15.13 | < $2 \times 10^{-16}$ |
| **Days post feeding** | -0.15 | 0.060 | -2.57 | 0.010 |
| **strain(Palm)** | -4.50 | 0.85 | -5.31 | $1.07 \times 10^{-7}$ |
| **strain(Field)** | -8.64 | 1.57 | -5.50 | $3.88 \times 10^{-8}$ |
| ***Wolbachia*(None)** | 0.17 | 0.21 | 0.82 | 0.41 |
| ***Wolbachia*(wPip)** | -0.044 | 0.19 | -0.23 | 0.82 |
| **Days post feeding*Palm** | 0.38 | 0.077 | 4.92 | $8.56 \times 10^{-7}$ |
| **Days post feeding*Field** | 1.01 | 0.15 | 6.78 | $1.20 \times 10^{-11}$ |

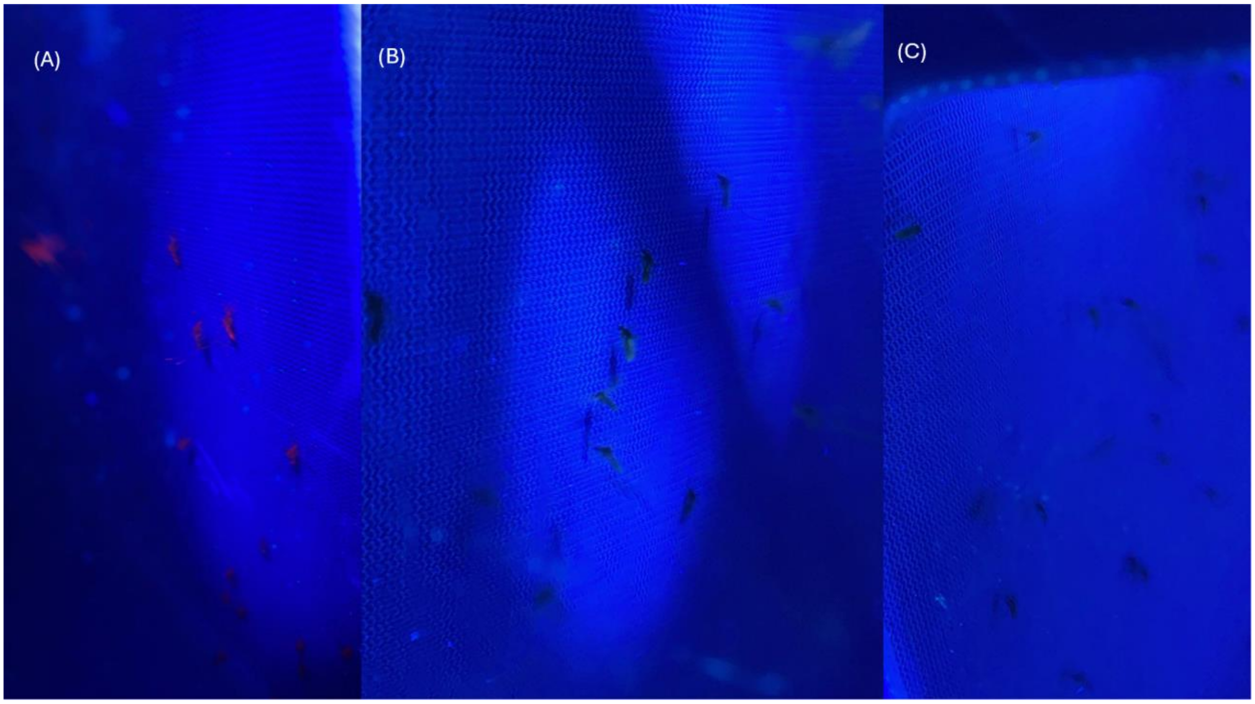


**Figure S1. Photographs under ultraviolet light of mosquitoes sprayed (A) red, (b) green, or unsprayed (c).**

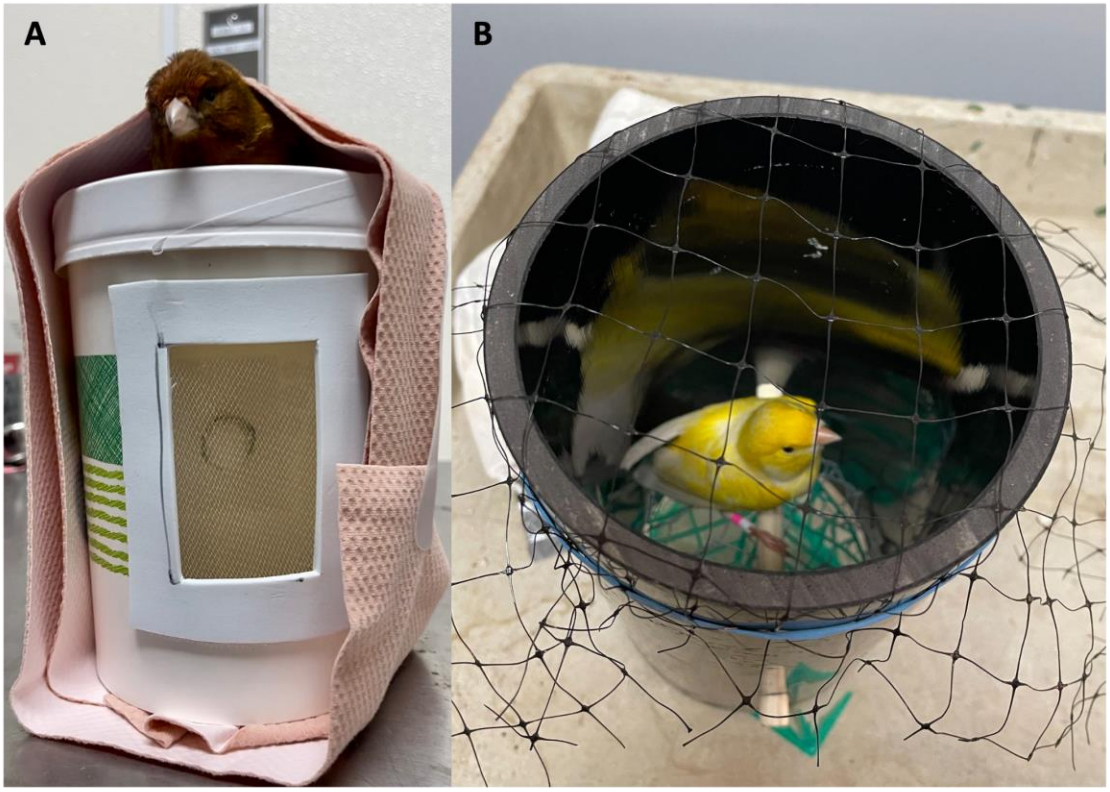


**Figure S2. Mosquito feeding containers with A) restrained and B) unrestrained canaries.** Feeding with both methods occurred in a room at 24 °C with the lights turned off. (A) The canary was restrained with a flexible athletic bandage on top of a 16-oz lidded paper container. The canary's legs are pulled through holes in the lid so they are inside the container where mosquitoes can feed on them. Mosquitoes are placed into the container through an opening at the back using a mouth aspirator (Model 612, John W. Hock), which was then closed with a cotton ball to keep mosquitos inside. A mesh screen at the front of the container allowed monitoring of the feeding process. The container was placed inside a BugDorm mosquito cage (4S3030, W30 x D30 x H30cm). (B) The canary was unrestrained and sat on a wooden perch inside a vertical PVC cylinder (10 cm diameter x 30 cm height). The cylinder was elevated on a wire platform and capped with plastic netting. This allowed mosquitoes to access birds from above and below and prevented the bird from flying. The cylinder was then placed inside a mosquito cage (BugDorm-4S3030, W30 x D30 x H30cm) that was subsequently filled with mosquitoes.

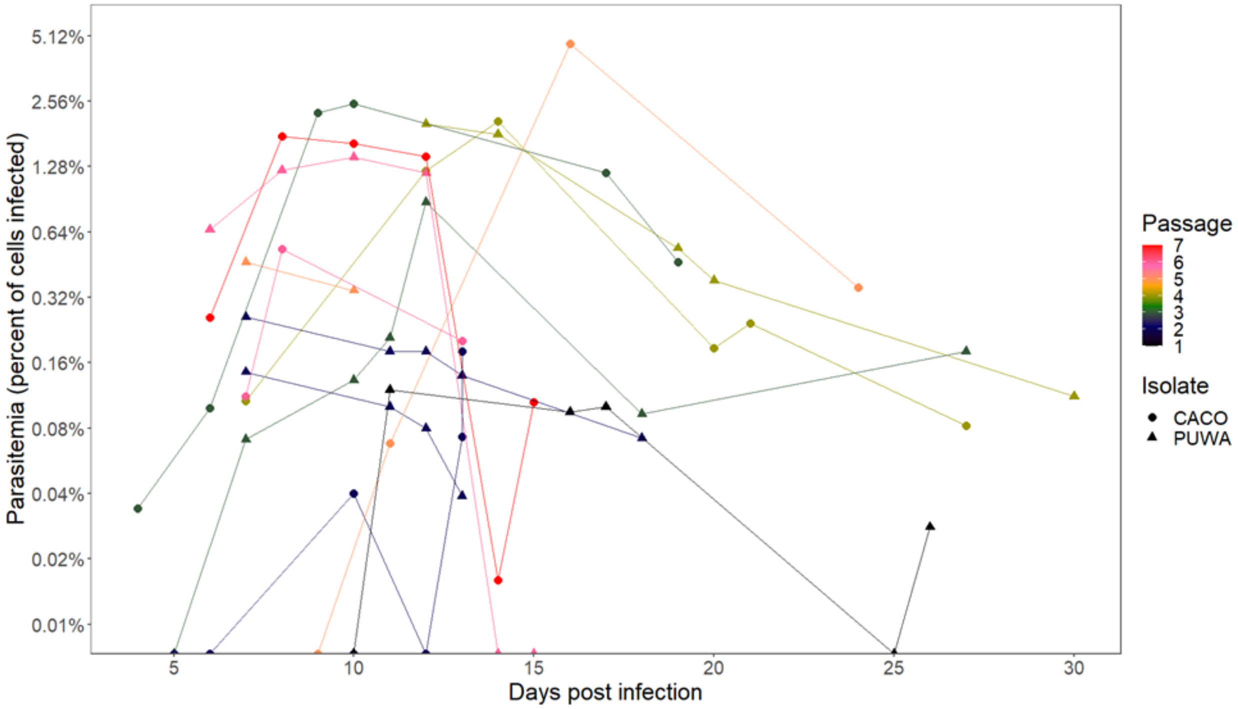


**Figure S3. Parasitemias (percent of red blood cells infected) over time in eighteen domestic canaries infected with *Plasmodium relictum* GRW4.** Points show parasitemia estimates measured from thin blood smears using microscopy. Colors show the passage number for the malaria isolate and shape shows the isolate (CACO - Captain Cook; PUWA - Pu’u Wa'awa'a).

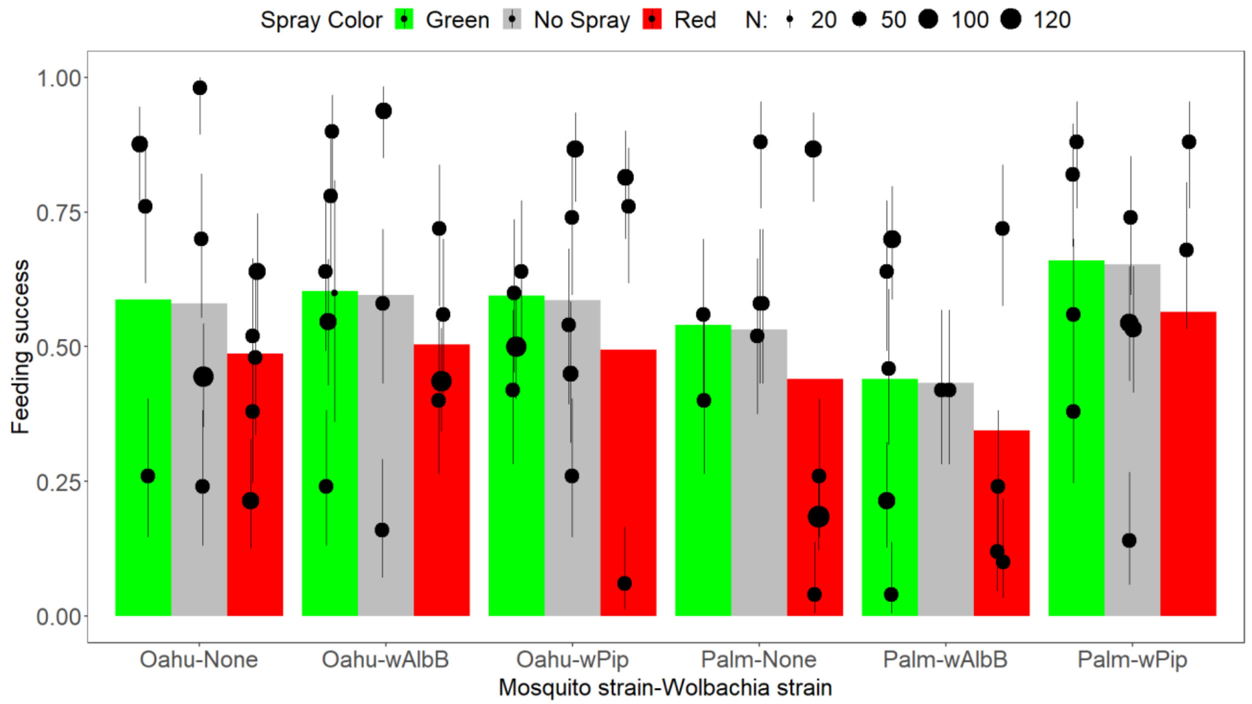


**Figure S4. Feeding success for two strains of mosquitoes (Oahu and Palmyra (Palm)), each with two strains of *Wolbachia* (*w*AlbB, *w*Pip) or no *Wolbachia* (None), with two colors of spray (green or red) or no spray.** Points show values from individual experiments (with binomial 95% CIs) and the size of the points shows the sample size of mosquitoes (range 20-130). Bars show the fitted model for each of six strain-*Wolbachia* combinations, with the color indicating the spray color.

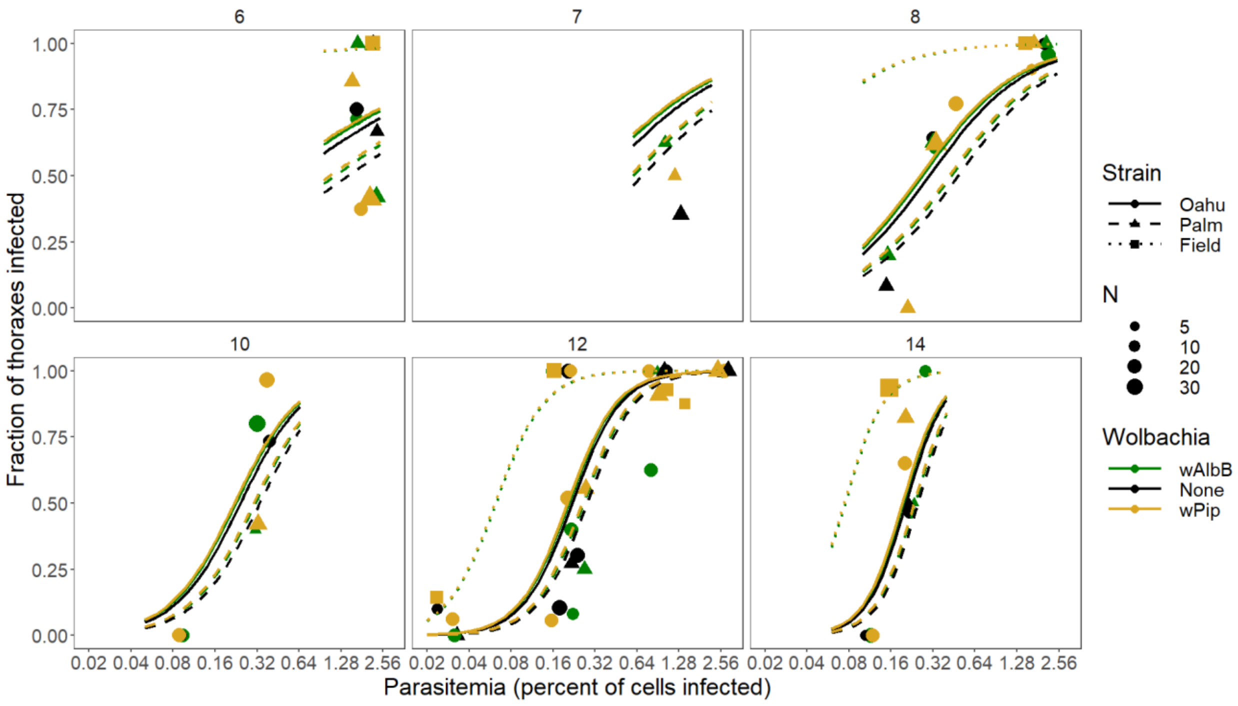


**Figure S5. Fraction of thoraxes infected plotted against the parasitemia (percent of red blood cells infected) of the bird the mosquitoes fed upon (on a $\log_2$ scale).** The color shows the *Wolbachia* type in the mosquitoes (*w*AlbB, *w*Pip, or None), the symbol and line type show the mosquito strain (colonized Oahu or Palmyra mosquitoes, or Field-type mosquitoes from Hawaii island), the different panels show the days post-feeding when the mosquitoes were dissected (6-14 days), and the size of the points shows the sample size (range 4-39). The lines show the fitted model for each of the three *Wolbachia* types, which are often on top of each other and difficult to distinguish because there is no statistical support for differences among *Wolbachia* types. Points have been slightly jittered along the x-axis to aid in visualization. Fitted lines are only shown on panels where there was data for that day post-feeding for that mosquito strain (i.e., there were no Field-type mosquitoes tested on days 7 and 10 so no fitted lines are shown).

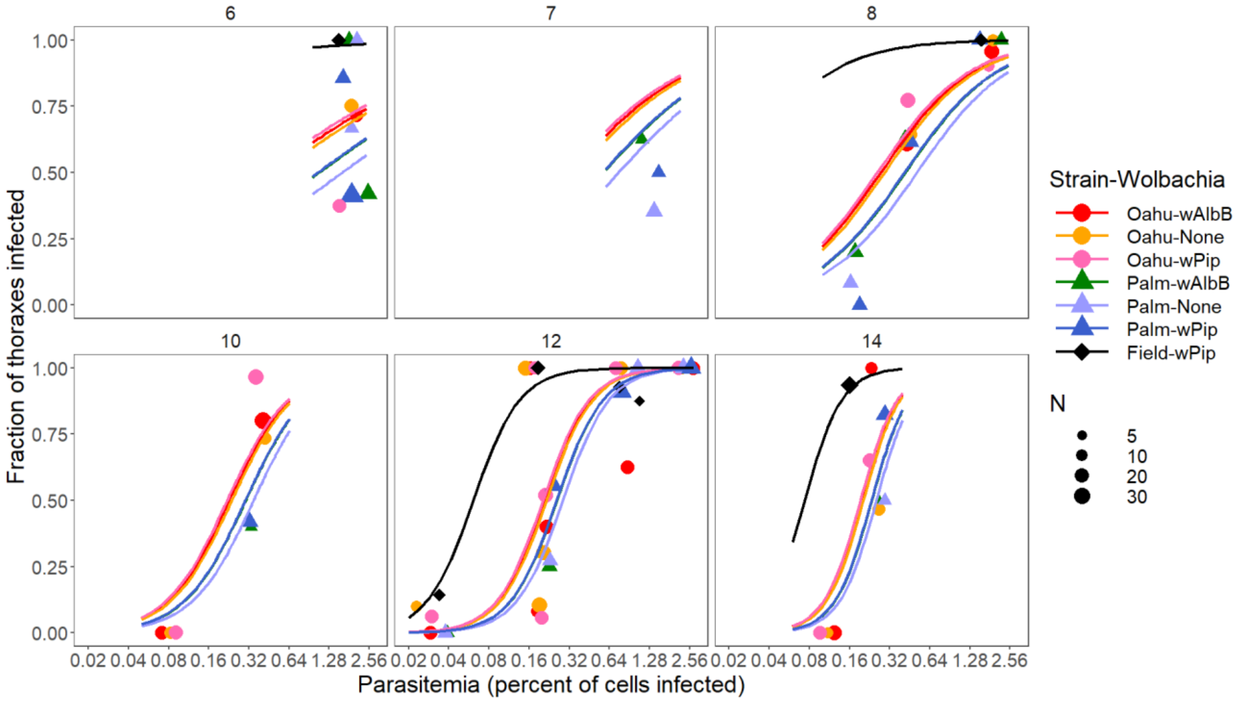


**Figure S6. Fraction of thoraxes infected plotted against the parasitemia of the bird the mosquitoes fed on (on a log$_2$ scale).** Color shows the mosquito strain (Oahu, Palm, Field) and *Wolbachia* strain (*w*Pip, *w*AlbB, none) pair, the different panels show the days post feeding when the mosquitoes were dissected (6-14), and the size of the points shows the sample size (range 4-39). The lines show the fitted model for each strain-*Wolbachia* pair; pairwise comparisons show differences among the three mosquito strains but no differences between *Wolbachia* strains within each mosquito strain. Points have been jittered along the x-axis to aid in visualization. Fitted lines are only shown on panels where there was data for that day post-feeding for that mosquito strain (i.e., there were no Field-type mosquitoes tested on days 7 and 10 so no fitted lines are shown).

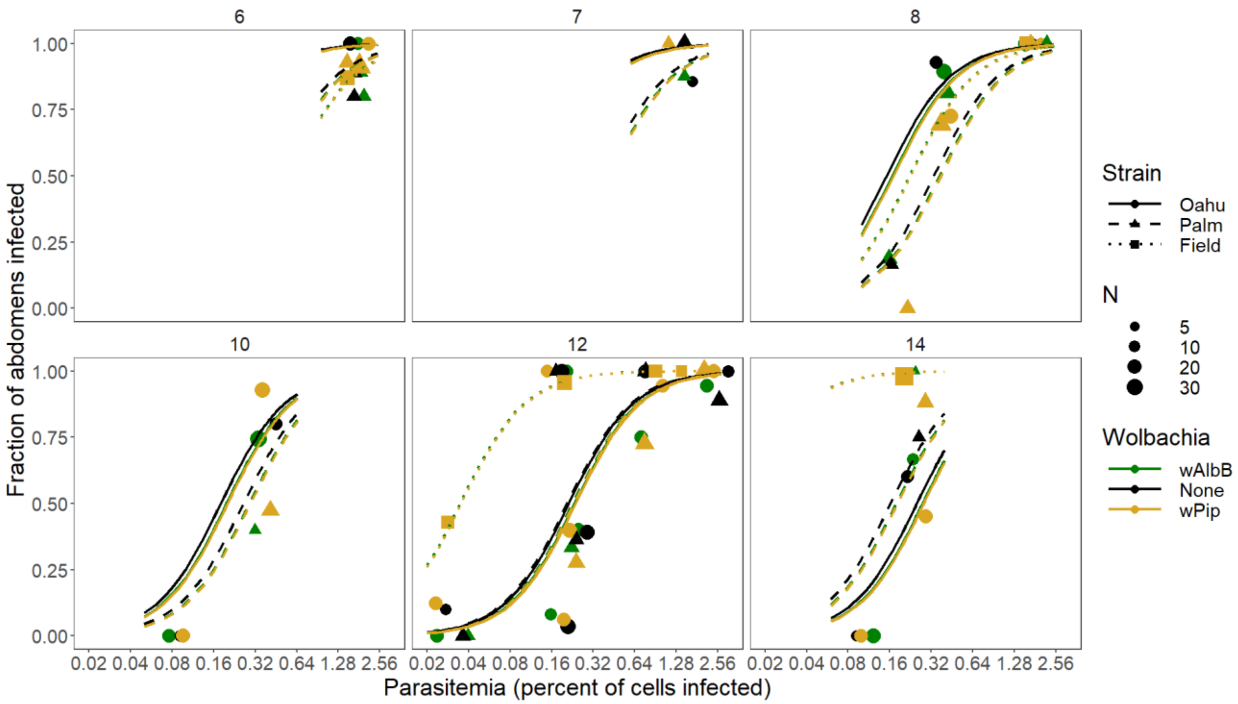


**Figure S7. Fraction of abdomens infected plotted against the parasitemia (percent of red blood cells infected) of the bird the mosquitoes fed upon (on a log$_2$ scale).** The color shows the *Wolbachia* type in the mosquitoes (*w*AlbB, *w*Pip, or None), the symbol and line type show the mosquito strain (colonized Oahu or Palmyra mosquitoes, or Field-type mosquitoes from Hawaii island), the different panels show the days post-feeding when the mosquitoes were dissected (6-14 days), and the size of the points shows the sample size (range 4-39). The lines show the fitted model for each of the three *Wolbachia* types, which are often on top of each other and difficult to distinguish because there is no statistical support for differences among *Wolbachia* types. Points have been slightly jittered along the x-axis to aid in visualization. Fitted lines are only shown on panels where there was data for that day post-feeding for that mosquito strain (i.e., there were no Field-type mosquitoes tested on days 7 and 10 so no fitted lines are shown).

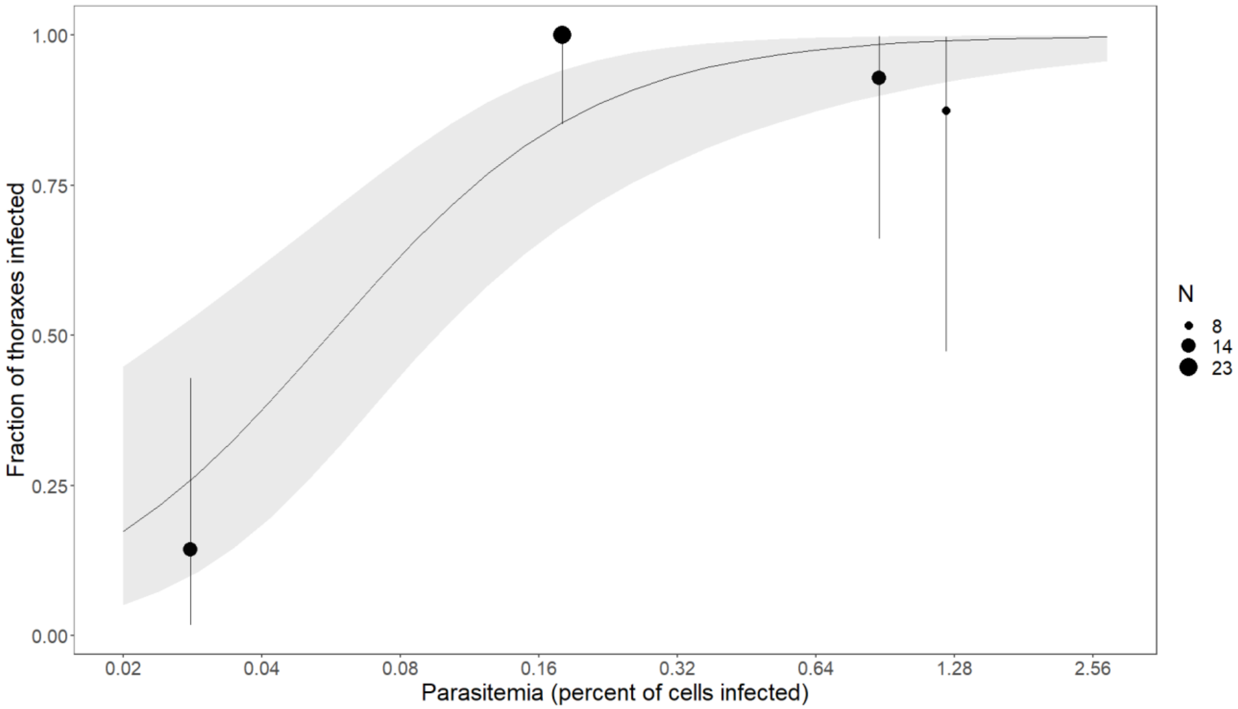


**Figure S8. The fraction of thoraxes infected in Field *C. quinquefasciatus* twelve days after feeding, plotted against parasitemia (percent of red blood cells infected).** Points show the data and 95% CI binomial intervals, and the line and ribbon shows the fitted relationship (Prevalence = logit(4.37 + 1.52*log(Percent parasitemia); slope SE = 0.39; Z-value = 3.89; P = 0.0001). This 95% CI of this slope (0.76, 2.28) is much lower and does not contain the slope of log parasitemia in a fit of the dataset without the Field strain (slope on day 12: -1.38+0.337*12 = 2.66). This is similar to the estimated slope for the full dataset, which was primarily composed of data from the inbred mosquito strains (Table S5: slope on day 12: -1.22+12*0.310 = 2.50).
